# Dissociable electrophysiological measures of natural language processing reveal differences in speech comprehension strategy in healthy ageing

**DOI:** 10.1101/2020.04.17.046201

**Authors:** Michael P. Broderick, Giovanni M. Di Liberto, Andrew J. Anderson, Adrià Rofes, Edmund C. Lalor

**Affiliations:** School of Engineering, Trinity Centre for Bioengineering and Trinity College Institute of Neuroscience, Trinity College Dublin, Dublin 2, Ireland; Laboratoire des systèmes perceptifs, Département d’études cognitives, École normale supérieure, PSL University, CNRS, 75005 Paris, France; Department of Biomedical Engineering, University of Rochester, Rochester, NY 14627, USA; Department of Neuroscience, and Del Monte Institute for Neuroscience, University of Rochester, Rochester, NY 14627, USA; Department of Neurolinguistics and Language Development, University of Groningen, Oude Kijk in Het Jatstraat 26, 9712EK Groningen, The Netherlands

**Keywords:** Ageing, N400, EEG, Language, Prediction, Computational Language modelling

## Abstract

Healthy ageing leads to changes in the brain that impact upon sensory and cognitive processing. It is not fully clear how these changes affect the processing of everyday spoken language. Prediction is thought to play an important role in language comprehension, where information about upcoming words is pre-activated across multiple representational levels. However, evidence from electrophysiology suggests differences in how older and younger adults use context-based predictions, particularly at the level of semantic representation. We investigate these differences during natural speech comprehension by presenting older and younger subjects with continuous, narrative speech while recording their electroencephalogram. We use linear regression to test how distinct computational measures of 1) semantic dissimilarity and 2) lexical surprisal are processed in the brains of both groups. Our results reveal dissociable neural correlates of these two measures that suggest differences in how younger and older adults successfully comprehend speech. Specifically, our results suggest that, while younger and older subjects both employ context-based lexical predictions, older subjects are significantly less likely to pre-activate the semantic features relating to upcoming words. Furthermore, across our group of older adults, we show that the weaker the neural signature of this semantic pre-activation mechanism, the lower a subject’s semantic verbal fluency score. We interpret these findings as prediction playing a generally reduced role at a semantic level in the brains of older listeners during speech comprehension and that these changes may be part of an overall strategy to successfully comprehend speech with reduced cognitive resources.

## Introduction

Healthy ageing is accompanied by a myriad of sensory and cognitive changes. This includes a decline in working memory (1) and episodic memory (2) as well as hearing loss (3) and a slowing in processing across cognitive domains (4). It is likely that changes in all of these faculties play into the reported extra difficulties that older adults experience in trying to follow everyday conversational speech, especially in challenging listening environments (5–7). While impaired hearing certainly plays a role in these difficulties (8, 9), it is also clear that “normal” hearing is not enough to guarantee good speech comprehension in everyday communication (10). But precisely how age-related changes in memory and speed of processing impact upon speech comprehension is less clear. In general, older adults show a relatively preserved language system and semantic memory (11). However, neuroimaging studies indicate key changes that occur with age that could impact the processing of speech at higher linguistic levels (12). As such, to better account for the communication challenges faced by older people, it is essential that we better understand how linguistic processing at these levels might differ between young and old.

One way in which researchers have explored age-related differences in the neurophysiology of language is via the N400 component of the event-related potential (ERP) (13, 14). The N400 is a centroparietal negativity that is elicited 200-600ms after word-onset and is strongest for words that are incongruent with their preceding context (e.g., “I take my coffee with cream and <u>*salt*</u>”). Several contrasting theories have been advanced to account for the N400. These include suggestions that the N400 reflects analysis of the low-level (e.g., orthographic or phonological) attributes of the unexpected (read or heard) word before that word is actually recognized (15); that it represents the process of accessing the semantic meaning of the word (16); or that it represents the process of incorporating the meaning of the word into its preceding context (17). One idea that has the potential to unify several of these competing theories is that the N400 reflects the stimulus induced change in a multimodal neural network, wherein an implicit and probabilistic representation of sentence meaning is held (14, 18). Importantly, the state of this internal network can be shaped by predictions, such that information can be partially or fully activated before the arrival of bottom-up input. This idea relies on the suggestion that listeners process speech predictively. In particular, it has been suggested that listeners use context to predictively pre-activate information at multiple representational levels during language comprehension (19). At a *lexical* surface level, this could include the activation of representations of word identity (20, 21), whereas a higher *semantic* level relates to the activation of an upcoming word’s semantic features (22). This is illustrated by an example sentence in Fig. 1. Indeed, from this perspective, it is conceivable that prediction at multiple representational levels could concurrently contribute to the N400 component.

**Fig 1:**
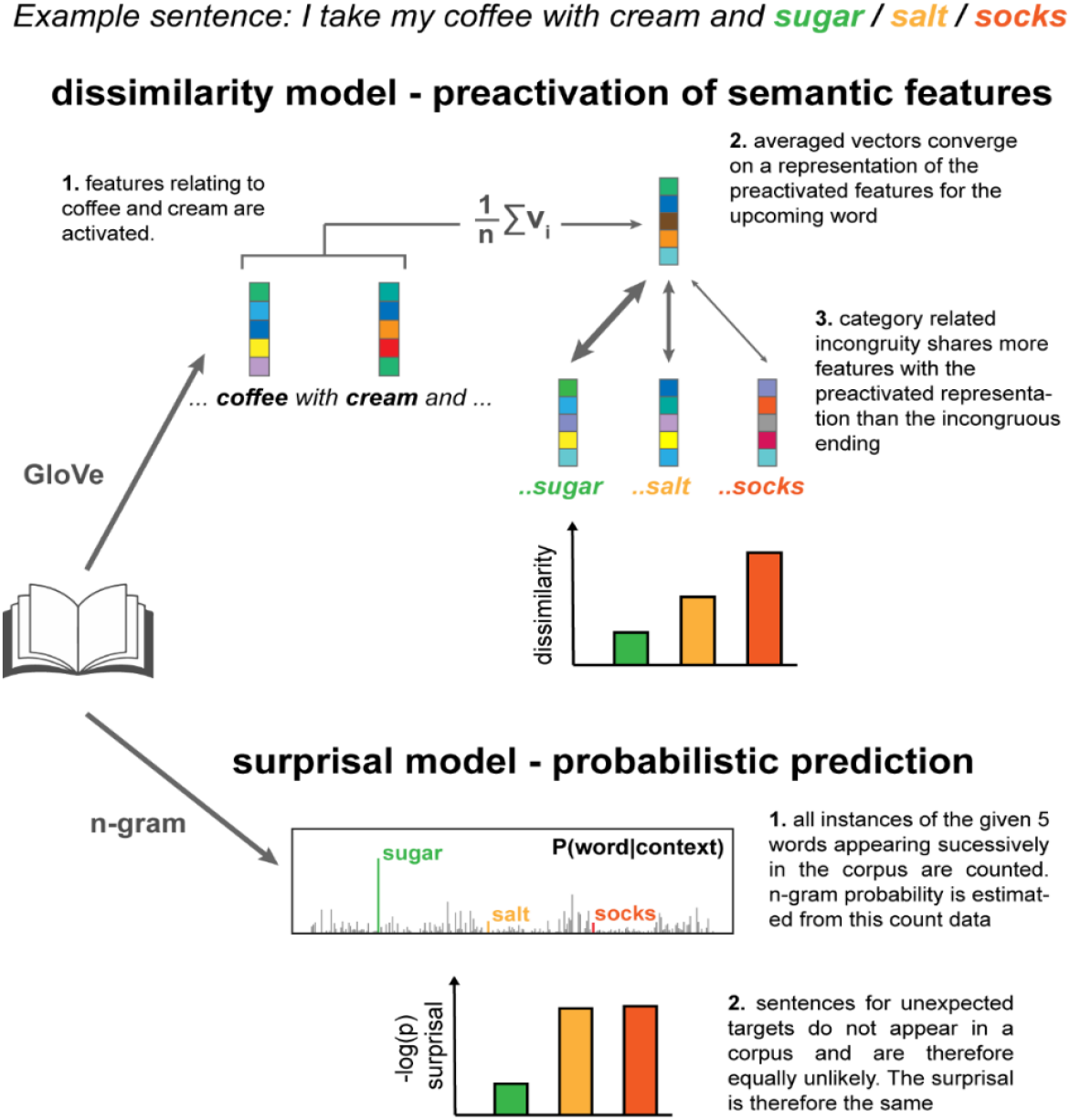
Computational models of predictive processing at lexical and semantic level. To illustrate the idea of prediction operating across multiple representational levels, consider the sentence *I take my coffee with cream and…* which ends with either an expected completion (*sugar),* an unexpected but semantically related completion (*salt*) or an unexpected and semantically unrelated completion (*socks*). At a lexical level, *salt* is unexpected because it is extremely rare that this sequence of words is heard or read. Processing of this word is therefore assumed to be no different from the processing of other unexpected words (i.e. *socks*). Conversely, at a higher *semantic* level, *salt* is relatively more likely, because *sugar* and *salt* share common features, both being powders and condiments; edible; white etc. We used two models of lexical surprisal and semantic dissimilarity to disentangle the contributions of prediction at lexical and semantic levels, respectively. **Top:** For the semantic dissimilarity model, vector representations of previous words in the sentence are averaged to form an estimation of the event context. The latent semantic features of the averaged vector converge on a representation similar to the predicted target “sugar” which, consequently, is more similar to words from the same category (e.g. “salt”) than different categories (e.g. “socks”). **Bottom:** Conversely, the lexical surprisal model does not distinguish between unexpected words based on their semantic category as it only reflects the probability of encountering either sequence of words in the training corpus, which is either rare or non-existent.

While the N400 component has been useful in studying age-related differences in the neurophysiology of language, the lack of consensus over what it reflects has complicated the interpretation of results in this area. The idea that the N400 might reflect differences in how younger and older adults use context-based predictions is evident in results from previous studies, particularly at the level of semantic representation (23). However, such results have been interpreted as older adults relying less on prediction *in general* during language comprehension, instead having responses that pattern more with plausibility ratings (12, 23). An alternative explanation for these differences is that ageing affects predictive processing at specific, semantic levels of representation rather than across all representational levels. This explanation is more consistent results from eye-tracking studies where it is believed that older adults rely more heavily on context-based probabilistic predictions (24, 25). But, again, the notion that predictive processes at multiple representational levels might contribute concurrently to the N400 and how this might be affected by ageing has received less attention.

In this study, we test whether prediction at distinct linguistic levels is differentially affected by ageing. To do this, we leverage a recent experimental framework (26) to isolate neural correlates of prediction from these different levels in younger and older adults using natural, continuous speech and modern context-based language modelling. This approach includes the variations in predictability at different levels that come with natural speech and allows for the derivation of interpretable neural correlates of different aspects of predictive language processing according to the language models used in analysing the neural data (27). Furthermore, the use of natural speech material adds to the ecological validity of observed effects and is less taxing on the attention of participating subjects than experiments involving artificially constructed sentences. This is important for reducing the potential confound of different levels of attentional engagement between older and younger subjects.

We exploit a recent modelling framework (Fig. 1) to tease apart neural correlates of predictive processing at the lexical and semantic level. To model predictive processing at the lexical level, we estimated 5-gram surprisal: an information theoretic measure of the inverse of the probability of encountering a word, given the ordered sequence of the 4 preceding words (28). In short, high lexical surprisal values arise from improbable word sequences. To model predictive processing at the semantic level, we exploited a popular distributional semantic modelling approach (29) that approximates word meaning, using numeric vectors of values reflecting how often each word co-occurred with other words across a large body of text. Distributional modelling approaches like this support the construction of conceptual knowledge hierarchies, e.g., a dragonfly is an insect is an animal (30), and would be expected to capture similarities between words belonging to similar categories, such as sugar and salt, and their difference to, say, socks. Thus, semantic dissimilarity would predict a greater N400 for “I take my coffee with cream and *socks*”, than for “I take my coffee with cream and *salt”* (31). In contrast, a 5-gram surprisal model would likely regard *salt* and *socks* as being equally unexpected at the lexical level because their occurrences are both, presumably, non-existent in a text corpus. So, using this model, one might not expect to see much difference in the N400 for *salt* vs *socks*. (Fig 1).

Given the differences in how these models operate, we hypothesized that they could be used to dissociate predictive processes at lexical and semantic representational levels in terms of how they contribute to the N400. Additionally, based on previous N400 literature (23), we hypothesized that older participants would show a specific detrimental effect in their predictive processing at the semantic level, and that this effect would correlate with behavioural measures of verbal fluency.

## Results

Two groups of 19 older (55-77 years, mean=63.9) and 19 younger (19-38 years, mean=26.8) subjects listened to the same 12-minute long excerpt of narrative speech while their electrophysiological (EEG) signal was recorded. The neural tracking of lexical and semantic information in the speech signal was assessed using a lagged linear regression. Specifically, this method models neural responses to speech by estimating a temporal filter that optimally describes how the brain transforms a speech feature of interest into the corresponding recorded neural signal. The filter, known as the temporal response function (TRF), consists of learned weights at each recorded channel for a series of specified time-lags. The TRF has typically been used to measure the cortical tracking of acoustic and linguistic properties continuous speech (32–34). However, recent approaches using this method have sought to represent continuous speech beyond its low-level acoustic features, in terms of its higher-level lexical-semantic properties (35, 36). For our speech stimulus, we estimated the lexical surprisal and semantic dissimilarity values for each content word and modelled neural responses to these features with the TRF.

Lexical surprisal is a measure derived from the probability of encountering 5 words (symbols) in a particular sequence, in a steady stream of words. The surprisal estimate itself captures nothing about what words mean, in the sense that it supplies no measure of whether *cats* and *tigers* are categorically similar, or whether either are domestic. However, word symbol sequences do in part reflect the structure of events in the real world (*cats* chase *mice*). They also reflect grammatical constraints on permissible symbol sequences (“*The jumped the a cat”* is nonsensical). In contrast, the semantic dissimilarity measure captures differences in the semantic categories that words belong to, and in the contexts that words appear in (*cats* and *tigers* are both felines, but *tigers* rarely occur in domestic contexts). Similar measures have been used to successfully model observed N400 phenomenon relating to semantic categories (31). Dissimilarity between a word and its preceding context was computed by 1 minus the Pearson’s correlation between the current word vector and the averaged vectors of all previous content words in the same sentence (36). We found that lexical surprisal and semantic dissimilarity were only weakly correlated (Pearson’s R = 0.14, p = 1.2 × 10^−5^, n = 916, Fig. S1A), indicating that they captured distinct features of the stimulus. Figure S1B provides example sentences with lexical surprisal and semantic dissimilarity values of the final word.

To fit the TRF, lexical surprisal and semantic dissimilarity were represented as vectors of impulses at the onset of each content word whose heights were scaled according to their surprisal or dissimilarity value. We regressed these vectors simultaneously to the recorded EEG signal of each individual participant. This produced separate TRF weights for surprisal and dissimilarity. Figure 2A shows the surprisal TRF weights for older and younger groups at midline parietal electrodes with scalp weight topographies at selected time windows (300ms, 400ms and 500ms; window width of 50ms) plotted above and below. Both groups show a prominent negative component, characteristic of the classic N400 ERP. We found that the latency of this component was significantly delayed by 74ms in the older group (T = 3.5, p < 0.005, 2 sample t-test, Cohen’s d = 1.13) and observed a correlation between age and response peak latency within the older group (Pearson’s r= 0.46, p = 0.047). Figure 2B shows the TRF weights for the semantic dissimilarity feature in older and younger subjects. Younger subjects showed comparable responses for dissimilarity and surprisal feature weights. In contrast, dissimilarity weights were significantly weaker than surprisal weights for older subjects (p<0.05 running paired t-test, FDR corrected).

**Fig 2:**
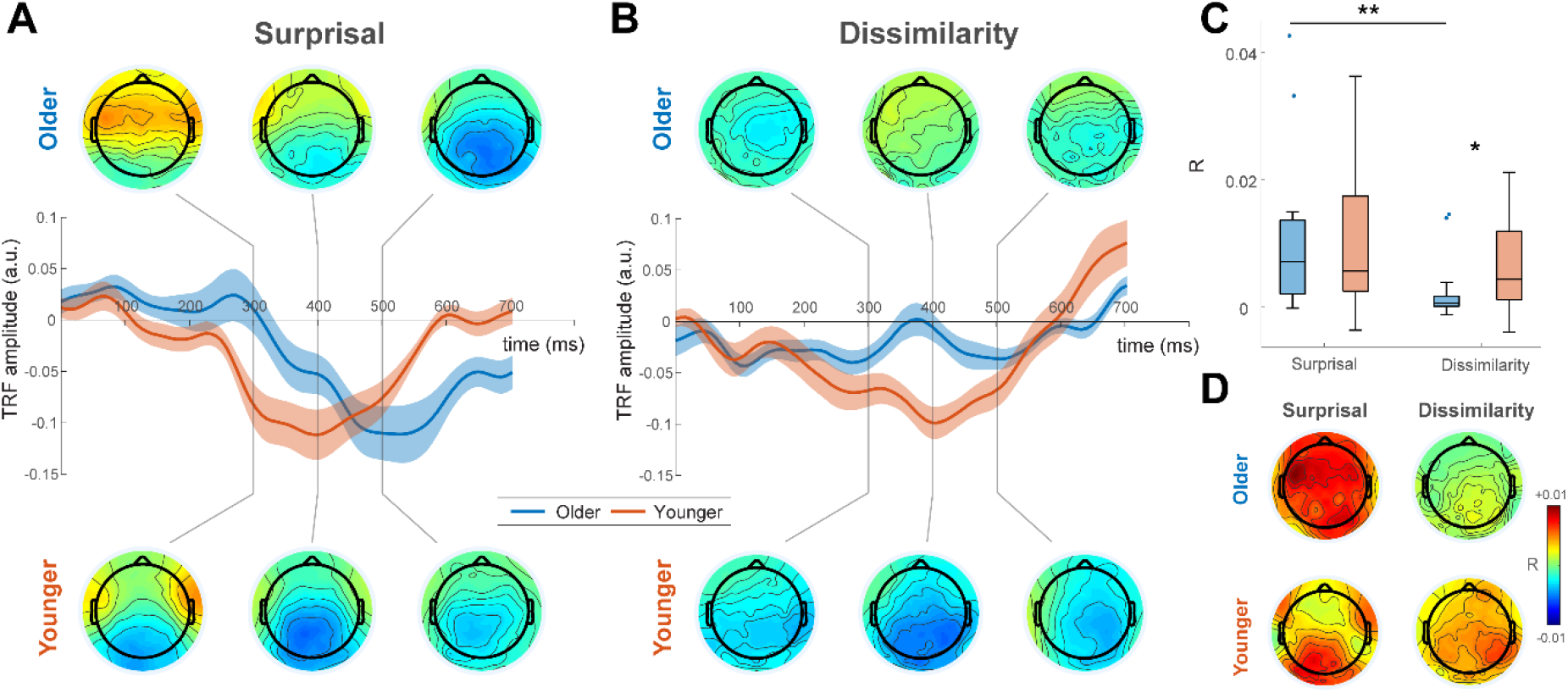
TRF weights and prediction accuracies. **A)** Lexical surprisal TRF weights averaged over parietal electrodes and across older (blue) and younger (red) subjects. Shaded areas show s.e.m. across subjects. N400 components are seen in the TRF weights at later time-lags for both groups and the peak latency of this component is significantly delayed for the older group. **B)** In contrast, semantic dissimilarity TRF weights are significantly weaker at later time lags for the older group compared to the younger group. **C)** Trained TRFs were used to predict unseen EEG in a cross-validation procedure. Consistent with the feature weights of the TRF, there were no significant differences between the prediction accuracies for dissimilarity and surprisal over parietal channels for younger subjects. However, older subjects showed significantly higher prediction accuracy for surprisal compared to dissimilarity at these channels. **D)** Topographical plots of prediction accuracy for both age groups and both models.

The performance of a model is also assessed by its ability to predict unseen neural data. Employing a cross validation procedure, we used each subject’s trained TRF model to predict their held-out EEG data. To test the predictive strength of surprisal and dissimilarity individually, we compared prediction accuracy of the full model (including dissimilarity and surprisal) with 5 null models where either surprisal or dissimilarity values were randomly permuted. Figure 2C shows the prediction accuracy (r) of each feature relative to the average null model predictions over midline parietal channels for younger and older subjects. Figure 2D shows the topographical distribution of r values. For both groups, surprisal and dissimilarity could significantly predict EEG above this baseline (Younger subjects: p = 0.0005 and p = 0.0011 for dissimilarity and surprisal, respectively; Wilcoxon signed-rank test. Older subjects: p = 0.022 and p = 0.0002, for dissimilarity and surprisal, respectively; Wilcoxon signed-rank test). Consistent with the feature weights of the TRF, there were no significant differences between the prediction accuracies for dissimilarity and surprisal for younger subjects (p = 0.28, Wilcoxon signed-rank test). However, importantly, older subjects showed significantly higher prediction accuracy for surprisal compared to dissimilarity (p = 0.0048, Wilcoxon signed-rank test). Younger subjects also showed significantly higher prediction accuracy for dissimilarity than older subjects (p=0.033, Mann-Whitney U-test).

From these results it is evident that semantic dissimilarity is weaker at explaining the neural responses for older subjects compared to younger subjects. However, this difference in model performance could conceivably be due to the particular way in which we have computed semantic dissimilarity. For instance, it has been shown that older adults have reduced working memory capacity (1), and thus for older adults it may be more appropriate to compute dissimilarity using a smaller window of previous words. To safeguard against this possibility, we tested several semantic dissimilarity vectors, where dissimilarity was estimated by comparing a word with a fixed number of previous words. We used context window sizes of 3, 5, 7, 9 and 11 words. We found no differences between models with different context window sizes or the model with a sentence context window (p = 0.61 for the older group, p = 0.79, for the younger group, Kruskal-Wallis test), indicating that the difference in brain responses between younger and older participants was not the result of the selected parameters. Finally, we investigated the low-level acoustic tracking of the speech envelope in both groups to check if our results might be explained based on differences in low-level encoding of the speech signal. Consistent with previous reports (6, 37), we found significantly *stronger* tracking of the speech envelope in older adults (Fig. S2), suggesting that low-level acoustic processing does not explain the between-group differences we see in semantic dissimilarity.

Previous work has indicated that older adults with higher verbal fluency scores were more likely to engage predictive processes at the level of semantics, resulting in N400 response patterns that were more similar to their younger counterparts (23). On this basis, we tested whether semantic dissimilarity model performance could predict verbal fluency in our older subjects. Semantic verbal fluency (VF) scores were collected from all but 2 of the older subjects. We found that, when controlling for age, model prediction accuracies were positively correlated with these scores across subjects (Pearson’s R = 0.59, p < 0.02, Fig. 3). This reveals that semantic dissimilarity was more accurately modelled for older subjects with higher semantic fluency. The model accuracy for surprisal was not predictive of this measure (Pearson’s R = −0.11, p > 0.05).

**Fig 3:**
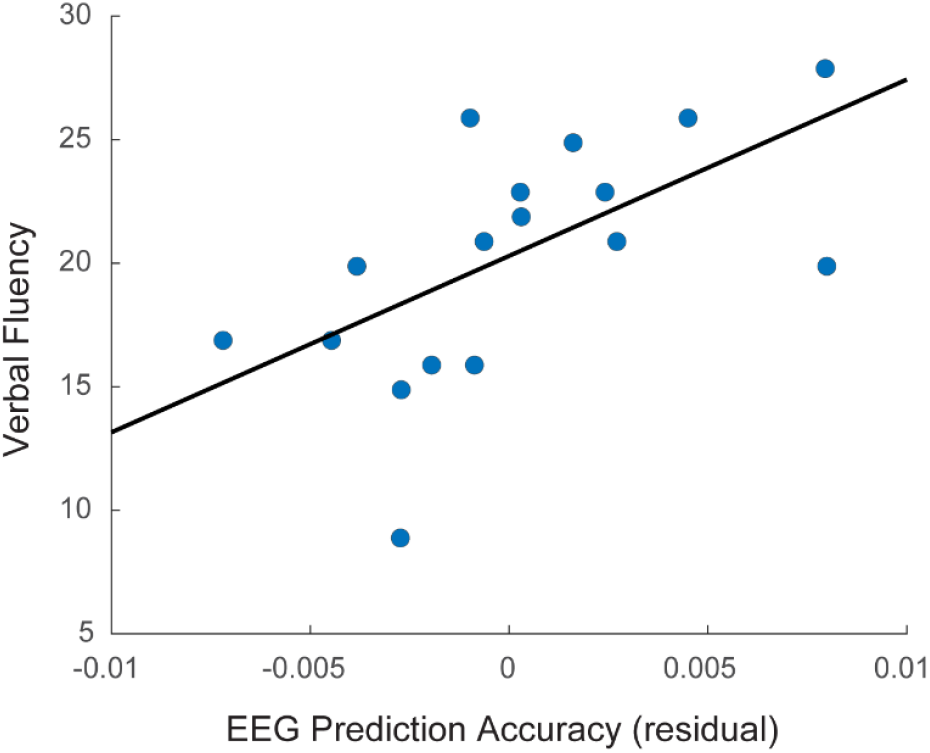
Within the older subjects, semantic category fluency was positively correlated with the semantic dissimilarity model performance when controlling for age (R = 0.59 p < 0.02)

## Discussion

The current article has revealed differences between younger and older adults’ electrophysiological responses to natural, narrative speech. In both young and old, a joint model capturing lexical surprisal and semantic dissimilarity produced N400 component responses in its temporal weights. While the lexical surprisal measure was robust in older adults, its peak negativity was delayed in a way that is similar to previous reports based on the N400 (13). In contrast, the semantic dissimilarity component was much reduced in the older subjects. We interpret this as evidence for two distinct contributions to the N400 that reflect predictive processing at the level of lexical and semantic representation. Furthermore, the pattern of results suggests that while older subjects maintain a robust ability to utilize lexical predictions during language processing, their ability to do so based on semantic representations appears to be impaired. Importantly, this interpretation was supported by the fact that the performance of the semantic dissimilarity model in older adults reflected a semantic behavioural measure of their categorical verbal fluency. These results extend basic scientific understanding of neurophysiological changes that accompany ageing and could have implications for research into naturalistic measures of brain health, as we discuss below.

The notion that our measures reflect predictive processing at different lexical-semantic levels fits with results from previous modelling studies on young adults. Surprisal and dissimilarity measures were jointly modelled on both EEG and fMRI responses in younger adults during sentence reading and narrative speech comprehension respectively (27). The measures both produced similar N400 responses in the sentence reading EEG data, as they have done for the younger adults in our study (Fig 2). However, the fMRI results provided evidence that distinct brain regions were involved in processing the two different aspects of the speech input. In particular, visual word form areas reflected surprisal (i.e., lexical processing) and areas of the semantic network (38) reflected dissimilarity (i.e., semantic processing). However, in that study, the distinct contributions of lexical and semantic processing were not dissociable in the N400 data because they were both strongly represented in younger adults. Our study goes beyond this, in 1) revealing that electrophysiological activity elicited in narrative speech comprehension reflects components of lexical surprisal and semantic dissimilarity; 2) demonstrating an age-related dissociation between the contribution of surprisal and dissimilarity, with the dissimilarity component being less pronounced in older people. Together, these findings provide convergent evidence that the N400 response reflects contributions from multiple processes relating to prediction at different levels of the linguistic processing hierarchy (19, 39).

Indeed, the idea that the N400 is affected by prediction at lexical and semantic levels fits with previous N400 ERP literature. N400 ERP responses in younger adults are enhanced for words that are not only unexpected in the context, but also belong to a different semantic category to the expected word (22). This is consistent with semantic features of upcoming words being predictively preactivated during comprehension. Importantly, the relative contribution of lexical and semantic components to the N400 appears to change with age. Specifically, in older adults there is less sensitivity to an unexpected word's semantic category, especially when the preceding context is highly constraining. These results are consistent with the idea of prediction playing a reduced role at a semantic level in the ageing brain (23, 40). However, whereas N400 ERP paradigms propose that different sub processes contribute to the N400 based on how evoked responses vary as a function of sentence-ending, our approach has dissociated lexical surprisal and semantic dissimilarity in neural responses to a continuous stretch of narrative speech. Furthermore, we have uncovered additional evidence that the semantic dissimilarity subprocess, specifically, yields a weaker response in older adulthood. Older adults show remarkably preserved language comprehension skills despite experiencing an overall decline in sensory and cognitive function (11). Changes in prediction at a semantic level may be part of a strategy to successfully comprehend speech with reduced cognitive resources (41). This possibly highlights the putative value of high-level predictions to support speech comprehension in noisy environments, when the input is corrupted and where older adults often struggle to comprehend. Future work, presenting speech at different levels of signal-to-noise ratio could help our understanding of such phenomena.

Two points we wish to further emphasize are firstly that the study was undertaken using a short, 12-minute segment of natural continuous speech stimulus. Previous research into the electrophysiological changes in language processing in the ageing brain have leveraged ERP-based experimental protocols that rely on experimenter-configured stimulus sets to enable contrasts between different stimulus conditions (e.g. congruent and incongruent sentence wordings). However, the ERP approach constrains the breadth of linguistic stimuli that can be investigated to the subset of sentences configured into matched experimental pairs. Additionally, the degree of ecological validity of results generated from bespoke ERP setups is unclear, because the experimental conditions are rarely experienced in everyday life. By examining electrophysiological responses elicited in audiobook comprehension, we have utilized a stimulus that is actually experienced in the wild, and the participant experiences an uninterrupted prolonged and cohesive discourse, that is likely to be more engaging than listening to disjoint experimental sentences. A second point we wish to emphasize is the current use of a predictive modelling approach. Previous ERP work on ageing tends to rely at least partially, on the experimenter to first configure stimulus categories to contrast and then populate them with sentences. It is challenging for a new experimenter to translate such an experiment to new stimuli, because they will have to manually select the new stimulus set. Conversely, the current modelling approach and the temporal response functions constructed in this experiment are portable and can be directly transferred to predict electrophysiological responses for entirely new narrative stimuli in new groups of young and old participants. However, we also wish to make explicit that the study of uncontrolled naturalistic stimuli, as opposed to tightly controlled N400 ERP sentence pairs, comes with its own limitations and presents a different rather than a “better” perspective on brain function. In the case at hand, it is unclear whether particular words and sentences in the audiobook played a key role in eliciting the electrophysiological response profiles and, if so, whether these critical sentences were the same for younger and older adults. In addition, the scope of experimentation is constrained by the content of the audiobook, which is not guaranteed to contain adequate variation in the linguistic factors of interest. This said, with the current set up, we have revealed a clear dissociation between lexical surprisal and semantic dissimilarity in the brains of older adults, thus establishing the general utility of using narrative speech to uncover age-related differences in high-level linguistic processing.

In conclusion, we have revealed neural correlates of language prediction relating to distinct measures of lexical surprisal and semantic dissimilarity. We show how one of these forms of prediction becomes less effective with age and patterns with behavioural cognitive measures, enabling us to predict an individual’s verbal semantic fluency from their neural data alone. These findings open new possibilities to study language impairment in the elderly and detect the onset of neurodegenerative disorders.

## Materials and Methods

### Participants

Data from 38 individuals (19 younger (6 female), age 19-38 years, M=26.8 years ± s.d. = 5 years; 19 older (12 female), age 55-77 years, M = 63.9 years ± s.d. = 6.7 years) was used in the study. Data from the younger subjects was collected in previous studies (34, 36) and in the current analysis only a portion of the dataset was used in order to match the data that was recorded for the older participants. Both studies were undertaken in accordance with the Declaration of Helsinki and were approved by the Ethics Committee of the School of Psychology at Trinity College Dublin. Each subject provided written informed consent. Subjects reported no history of hearing impairment or neurological disorder.

### Stimuli and Experimental procedure

The stimulus was an audio-book version of a popular mid-20th century American work of fiction (The Old Man and the Sea, Hemingway, 1952), read by a single male American speaker. The first 12 minutes of the audiobook was divided into 4 trials, each 3 minutes in duration. The average speech rate was 190 words/minute. The mean length of each content word was 334ms with standard deviation of 140ms. Trials were presented chronologically to the story with no repeated trials. All stimuli were presented monophonically at a sampling rate of 44.1 kHz using Sennheiser HD650 headphones and Presentation software from Neurobehavioural Systems. Testing was carried out in a dark, sound attenuated room and subjects were instructed to maintain visual fixation on a crosshair centred on the screen for the duration of each trial, and to minimise eye blinking and all other motor activities.

Older participants were additionally tested with 2 verbal fluency (VF) tasks. Letter verbal fluency was measured by asking participants to name as many words beginning with the letter ‘F’ as they could in 60 seconds. Similarly, semantic verbal fluency was measured by asking participants to name as many animals as they could in 60 seconds. Prior to the verbal fluency task, participants were screened using the Montreal Cognitive Assessment (MOCA). Individuals who scored below 25 out of 30 in this test did not complete this task and therefore VF measures from 2 of the 19 subjects were not obtained.

### EEG Acquisition and Preprocessing

128-channel EEG data were acquired at a rate of 512 Hz using an ActiveTwo system (BioSemi). Offline, the data were downsampled to 128Hz and bandpass filtered between 0.5 and 8Hz using a zero-phase shift Butterworth 4^th^ order filter. To identify channels with excessive noise, the standard deviation of the time series of each channel was compared with that of the surrounding channels. For each trial, a channel was identified as noisy if its standard deviation was more than 2.5 times the mean standard deviation of all other channels or less than the mean standard deviation of all other channels divided by 2.5. Channels contaminated by noise were recalculated by spline interpolating the surrounding clean channels. Data were then referenced to the average of the 2 mastoid channels.

Finally, we applied multiway canonical component analysis (MCCA) to denoise the data. MCCA is a technique that seeks to extract canonical components across subjects (42). Like CCA, which is applied to single subjects, it can be used to find linear components that are correlated between stimulus and response. However, rather than analysing the components directly, EEG can be denoised by projecting it to the overcomplete basis of canonical components, selecting a set of components and then projecting back to EEG space. We denoised each age-group separately, with the prior hypothesis that latency and morphology of the group responses would be different. For each group, we chose parameters of 40 principal components for the initial principal component analysis and then 110 canonical components. These chosen values were based on the parameters that were recommended for denoising speech related EEG (42); however, we tried several different parameter pairs and tested their effect on the prediction accuracy of EEG from the speech envelope. We found that the recommended parameters returned the optimal denoising for both groups as determined by prediction accuracy of EEG based on the speech envelope (Figure S2).

### Semantic dissimilarity and surprisal estimation

#### Semantic Dissimilarity

Distributed word embeddings were derived using GloVe (29). This method factorizes the word co-occurrence matrix of a large text corpus, in this case Common Crawl (https://commoncrawl.org/). The output is 300-dimensional vectors for each word, where each dimension can be thought to reflect some latent linguistic context. These word embeddings are used to calculate our semantic dissimilarity measure. This is an impulse vector, the same length as a presented trial, with impulses at the onset of each content word. The height of each impulse is 1 minus the Pearson’s correlation between that word’s vector and average of all preceding word vectors in the same sentence. Semantic dissimilarity values have a mean of 0.48 ± s.d. = 0.17.

#### Surprisal

Surprisal values were calculated using a Markov model trained on the same corpus as GloVe (common crawl). These models, commonly referred to as n-grams, estimate the conditional probability of the next word in a sequence given the previous *n−1* words. We applied a 5-gram model that was produced using interpolated modified Kneser-Ney smoothing (28, 43). Surprisal vectors were calculated as impulses at the onset of all words whose heights were scaled according to the negative log of a word’s 5-gram probability. Any impulses that were not common between dissimilarity and surprisal vectors were removed. Surprisal values were normalised to match the distribution of dissimilarity values with a mean of 0.48 ± s.d. = 0.16.

### Temporal Response Function

The forward encoding model or temporal response function (TRF) can be thought of as a filter that describes the brain’s linear mapping between continuous speech features, S(t), and continuous neural response, R(t).

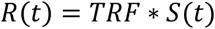

Where ‘*’ represents the convolution operator. The speech input can comprise of a single speech feature, i.e. univariate, or multiple speech features, i.e. multivariate. Each feature produces a set temporal weights for a series of specified time lags. TRF weights are estimated using ridge regression.

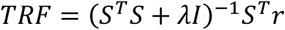

where λ is the regularization parameter that controls for overfitting. The models are trained and tested using a 4-fold cross-validation procedure. 3 of the 4 trials are used to train the TRF which predicts the EEG of the remaining trial, based on speech representation input. We train and test models based on the combined semantic dissimilarity and surprisal impulse vectors with the addition of an onset impulse vector with impulse height equal to the average dissimilarity and surprisal values across all words in the current trial. The onset vector acts as a nuisance regressor to capture variance relating to any acoustic onset responses. For testing, the prediction accuracy (R) of the model is calculated as the Pearson’s correlation between the predicted EEG and the actual EEG. A range of TRFs were constructed using different λ values between 0.1 and 1000. The λ value corresponding to the TRF that produced the highest EEG prediction accuracy, averaged across trials and channels, was selected as the regularisation parameter for all trials per subject.

To test directly how well each feature accurately captured neural activity for each subject we measured the model’s ability to predict EEG based on the true feature representation above null feature representations. Specifically, the heights of the impulses for the semantic models were randomly shuffled to produce permuted dissimilarity or permuted surprisal vectors. In the testing phase of the cross-validation procedure, a trained TRF would attempt to predict the neural response to the permuted features, while all other features remained constant. This was repeated for 5 permutations of each stimulus feature. Hence, prediction accuracy for semantic features refers to the prediction accuracy difference between true speech feature and the average of the 5 null speech feature representations.

In addition, we extracted properties of the model weights themselves. N400 peak latency was calculated automatically for each subject as the time lag with the lowest peak weight within a window of 200-600ms after a time-lag of zero. The response peak delay between groups was calculated as the difference between group averaged peak delays.

### Statistical Testing

For every statistical comparison, we first verified whether the distribution of the data violated normality and was outlier free. This was determined using the Anderson-Darling test for normality and 1.5 IQR criterion, respectively. We used parametric tests (t-test, paired t-test) for data which satisfied these constraints and non-parametric tests for data which violated them.

## Acknowledgements

We would like to thank Christian Buck for his help in providing 5-gram measures using Common Crawl.

