## Supplemental Information for "Dissociable electrophysiological measures of natural language processing reveal differences in speech comprehension strategy in healthy ageing"

### Supplementary Material

A

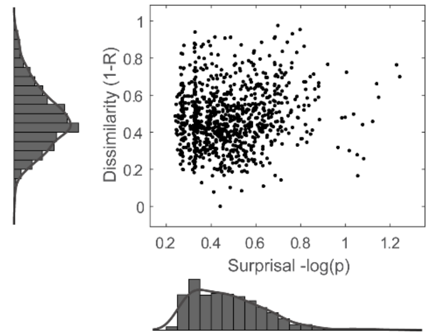


B

| Sentence | Dissimilarity | Surprisal |
|---|---|---|
| *"Those who had caught **sharks**"* | 0.36 | 0.8 |
| *"But none of these scars were **fresh**"* | 0.8 | 0.34 |
| *"But you are your father's and your **mother's**"* | 0.18 | 0.32 |
| *"I'll be back when I have the **sardines**"* | 0.93 | 0.76 |

**Fig. S1: Lexical surprisal and semantic dissimilarity values of individual words. A)** Marginal and joint distributions of the normalised surprisal and dissimilarity values for individual words. These values were weakly correlated (Pearson's R = 0.14, $p = 1.2 \times 10^{-5}$, n = 916). **B)** Example sentences with lexical surprisal and semantic dissimilarity values for the final word (indicated in bold). Four sentences with combinations of high and low surprisal and high and low dissimilarity word values were selected as examples.

##### Robust tracking of the speech envelope across age-groups

We conducted an additional analysis looking at the tracking of low-level acoustic properties of speech. The TRF approach has been commonly used to study the tracking of the broadband amplitude envelope of the speech signal (1, 2). Furthermore, it has been shown how this tracking is modulated by attention and speech intelligibility (3–6). We estimated TRFs using the speech envelope as input for both groups. Figure S2A shows the average weights for older and younger participants over bilateral temporal electrodes. Plotted above and below are the topographical distributions of these weights at selected time windows. For both groups, the temporal weights showed a morphology, consistent with previous studies (7). However, older subjects showed modulated peak responses relative to the younger group. P1, N1 and P2 components peaks were significantly larger for older subjects ($p < 0.05$ 2 sample t-test, false discovery rate (FDR) corrected). We also found age-related latency delays of 27ms, 10ms and 17ms for P1, N1 and P2 respectively ($p < 0.001$ 2-sample t-test). The accuracy of a model is also revealed in its ability to predict unseen EEG. Using a cross validation procedure, we used trained models to predict held out EEG data. Figure S2B shows the prediction accuracy (R) of the model for older and younger subjects at each channel (left) and for each subject averaged over 12 bilateral temporal electrodes (right). This shows an increase in prediction accuracy for older compared to younger participants (T = 3, p < 0.01 2-sample t-test, effect size (d') = 0.98). These findings are consistent with previous studies, showing stronger tracking of the speech envelope in older adults (8–10).

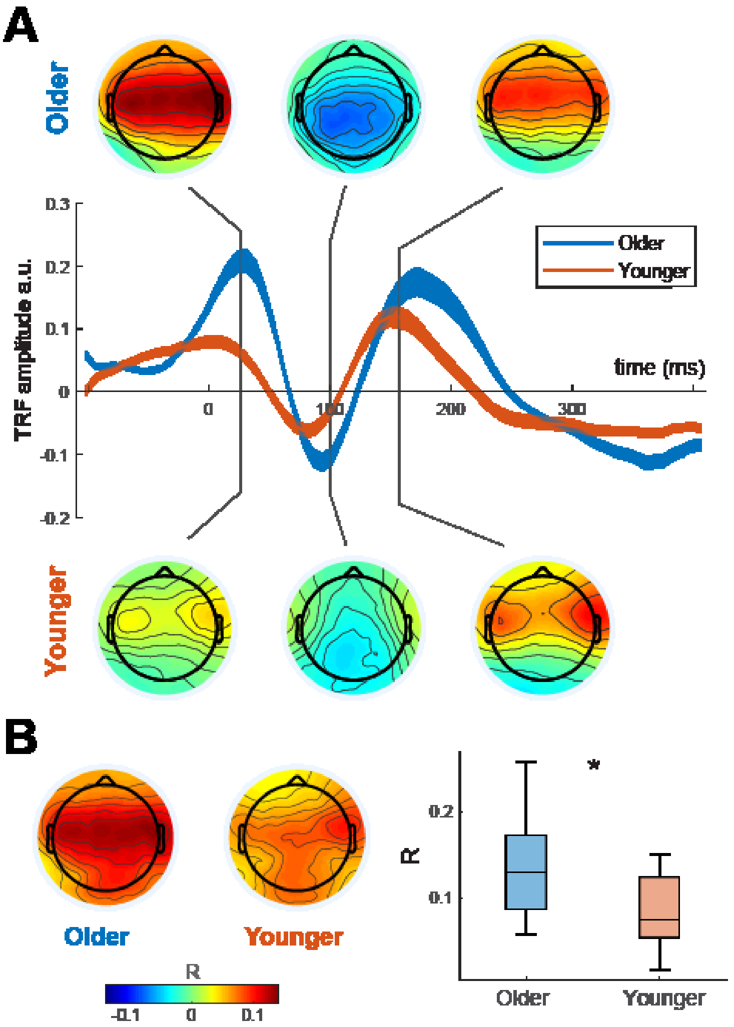


**Fig. S2: Modelling of acoustic processing in older and younger subjects. A)** A linear model trained on the speech envelope and recorded EEG signal reveals age-related differences in its temporal feature weights, averaged across subjects and over temporal electrodes. Topographies at selected time windows (~30ms, ~100ms and ~150ms) shows the same morphology of responses for each group. However, the response is stronger for older subjects. **B)** The trained models were used to predict unseen EEG in a cross-validation procedure. Older subjects had higher EEG prediction accuracy (R) compared to younger subjects.
